# Superoanterior Fasciculus (SAF): Novel fiber tract revealed by diffusion MRI fiber tractography

**DOI:** 10.1101/319863

**Authors:** Szabolcs David, Anneriet M. Heemskerk, Francesco Corrivetti, Michel Thiebaut de Schotten, Silvio Sarubbo, Laurent Petit, Max A. Viergever, Derek K. Jones, Emmanuel Mandonnet, Hubertus Axer, John Evans, Tomáš Paus, Alexander Leemans

## Abstract

Substantial progress in acquisition, processing, and analysis boosted the reliability of diffusion-weighted MRI and increased the accuracy of mapping white matter pathways with fiber tractography. Since the introduction of ‘region of interest’ (ROI) based virtual dissection by Conturo et al. in 1999, researchers have used tractography to identify white matter pathways, which faithfully represented previously known structures revealed by dyeing studies or post-mortem descriptions. The reconstructed streamlines are subjects of bundle-specific *in vivo* investigations to show differences between groups (e.g., comparing fractional anisotropy (FA) between patients and healthy controls) or to describe the relation between diffusion scalars and metrics of interest (e.g.: normal aging or changes due to learning). By applying a reverse strategy in using diffusion-weighted MRI tractography first, then supporting the findings with other techniques, we have identified a bilateral tract in the frontal cortex - the superoanterior fasciculus (SAF). The tract resembles the anterior shape of the cingulum bundle, but is located more frontally. To erase the chance that our findings are confounded by acquisition, processing or modeling artifacts, we analyzed a total of 421 subjects from four cohorts with different acquisition schemes and diverse processing pipelines. The findings were also completed with other non-MRI techniques, such as polarized light microscopy and dissection. Tractography results demonstrate a long pathway and are consistent among cohorts, while dissection indicates a series of U-shaped fibers connecting adjacent gyri. In conclusion, we hypothesize that these consecutive U-shaped fibers emerge to form a pathway, thereby resulting in a multicomponent bundle.

## 1 Introduction

Diffusion-weighted MRI (dMRI) based fiber tractography (FT) (Jeurissen et al., 2017) is widely used for investigating microstructural properties of white matter (WM) fiber bundles (Alexander et al., 2016) and for mapping structural connections of the human brain (Sotiropoulos and Zalesky, 2017; Wakana et al., 2004). Since the first endeavors of FT (Basser et al., 2000, 1994; Catani et al., 2002; Conturo et al., 1999; Jones et al., 1999; Mori et al., 1999), many studies have contributed to the improvement of data quality and the level of detail. Recent advances in MRI hardware and acquisition (Anderson, 2005; Andersson et al., 2016; Moeller et al., 2010; Setsompop et al., 2018, 2013; Sotiropoulos et al., 2013) have significantly improved dMRI data in terms of, for example, spatial resolution, angular resolution signal-to-noise ratio, and geometrical distortion reduction. Additionally, notable developments in data processing have reduced errors from the subjects’ head motion (Leemans and Jones, 2009), data outliers (Chang et al., 2005; Collier et al., 2015; Pannek et al., 2012; Tax et al., 2015), eddy currents (Andersson et al., 2016; Andersson and Sotiropoulos, 2015), EPI distortions (Andersson et al., 2003), Gibbs-ringing (Kellner et al., 2016; Perrone et al., 2015; Veraart et al., 2016), and other data artifacts, like drift of the diffusion signal (Vos et al., 2017). Besides the traditional diffusion tensor model, more sophisticated methods have been developed to resolve crossing fibers (Dell’Acqua et al., 2013; Jensen et al., 2016; Lin et al., 2003; Tax et al., 2014; Tournier et al., 2004; Tuch, 2004; Wu and Alexander, 2007). Together, all these improvements allow for more reliable FT results, thereby resolving complex fiber architecture, smaller branches of fiber bundles or minor fiber pathways (Tournier et al., 2008). Although dMRI based tractography is a promising technique, there are many well-known pitfalls and limitations in acquiring and analyzing dMRI data (Derek K. Jones et al., 2013; Jones and Cercignani, 2010; O’Donnell and Pasternak, 2015). dMRI is an indirect approach for measuring the underlying WM pathway properties, as it is an approximation of the average diffusion within a given voxel. Modeling the diffusion results in a proxy of main directions, and thus the exact architecture of the pathways cannot be determined unambiguously. The modeling errors can cause tracts to stop prematurely or jump from one WM structure to another, resulting in false negative (FN) or false positive (FP) connections. The strength of connectivity, when assessed via probabilistic FT, is unreliable due to its sensitivity to data quality (Mesri et al., 2016).

Despite recent efforts to increase the accuracy of FT by either pruning the tractograms using anatomical information (Roine et al., 2015; Smith et al., 2012) or by including microstructural information to disambiguate between pathways (Smith et al., 2013), the ISMRM Tractography Challenge in 2015 demonstrated that some data processing pipelines could result in large errors as the reconstructed tracts produced an average ratio of 1:4 in FP:FN connections and 45% in bundle overlap when compared to pre-defined, ground-truth streamlines (Maier-Hein et al., 2017).

In another study, the sensitivity (true connections) and specificity (avoidance of false connections) of dMRI tractography was investigated by combining tracer studies and high quality dMRI data (Thomas et al., 2014). This research showed a sensitivity-specificity trade-off, i.e. the number of true connections increased simultaneously with the number of false connections. Additionally, the anatomical accuracy of the tracts depended on the studied pathway, the acquisition and the processing pipeline. This stresses the fact, that comparing results across studies can suffer from the differences in acquisition settings and processing methods. Therefore, reconstructing fiber pathways from dMRI tractography should be performed with great care and additional support from other methods is welcome.

Nonetheless, being aware of the recognized issues, we present the description of a novel fiber tract, the superoanterior fasciculus (SAF), in the prefrontal lobe which – to the best of our knowledge – has not been documented before with dMRI based tractography with this level of detail. The tract is slightly curved as it follows the arc of the cingulum bundle and is located more frontally and above of the frontal part of the cingulum. Furthermore it can be found in both hemispheres.

Traditionally, the description of white matter fiber bundles went through the following course: tremendous amount of invasive evidence, for example histology (Dejerine and Dejerine-Klumpke, 1895) and dissection (Ludwig and Klingler, 1956) studies supported the presence of a structure, which later was confirmed via DTI based FT in the living human brain. Conturo et al. (Conturo et al., 1999) laid the foundations of ‘region of interest’ (ROI) based selection and analysis of tracts opening up a new era of tract based investigations from neuroscience (Lebel et al., 2008; Thiebaut de Schotten et al., 2012, 2011a) to clinical applications (Bartolomeo et al., 2007; Deprez et al., 2012; Thiebaut de Schotten et al., 2005). The access to large scale populations made even the construction of *in vivo* white matter tract atlases and guidelines feasible (Hua et al., 2008; Mori et al., 2005; Wakana et al., 2007). In this work, we also use specific ROIs along with constrained spherical deconvolution (CSD) (Tournier et al., 2007, 2004) based modeling of the diffusion MR signal to reconstruct the proposed structure.

To minimize the possibility that these findings are based on acquisition or processing artifacts, we used datasets from different projects: Human Connectome Project (HCP) (Glasser et al., 2013; Sotiropoulos et al., 2013; Van Essen et al., 2013), Avon Longitudinal Study of Parents and Children (ALSPAC) (Golding and ALSPAC Study Team, 2004), Multiple Acquisitions for Standardization of Structural Imaging Validation and Evaluation (MASSIVE) (Froeling et al., 2017), an in-house dataset (Jeurissen et al., 2011), and robust processing pipelines to show these trajectories. While the main focus of this research was based on dMRI, we complemented our findings with other non-MRI techniques, such as polarized light microscopy (PLI) (Axer, 2011) and dissection. Preliminary result of current work was presented at the ISMRM 23rd Annual Meeting and Exhibition 2015 in Toronto, Canada (Heemskerk et al., 2015).

## 2 Methods

In the methods section we describe the datasets that were used. Thereafter, the processing, tractography and analysis steps are explained. Finally, the PLI and dissection methods are described.

### 2.1 Diffusion MRI

#### 2.1.1 Datasets

Four different diffusion MRI datasets were used to investigate the possible existence of the tract among different platforms.

<u>Dataset 1:</u> The minimally processed dMRI data from the HCP S500 release was used. Every subject for whom all the 90 b = 3000 s/mm^2^ images were available with 18 b0 volumes, and were not listed among subjects with known anatomical anomalies or data quality issues were included in the analysis. The IDs of the excluded subjects are listed on the HCP wiki page (HCP wiki, 2018). The selection resulted in 409 healthy subjects. For a subset of 10 subjects we also analyzed the b=1000 s/mm^2^ and 2000 s/mm^2^ shells with both CSD and DTI models. <u>Dataset 2:</u> Ten dMRI datasets were used from the ALSPAC cohort, consisted of 60 diffusion gradients with b = 1200 s/mm^2^, and 2.4 mm isotropic voxel size as described by Golding and ALSPAC Study (Golding and ALSPAC Study Team, 2004). <u>Dataset 3:</u> The dataset is described in the work of Jeurissen et al. (Jeurissen et al., 2011). In summary, one subject, consisted of 60 diffusion directions, 2.4 mm isotropic voxel size (resampled to 1mm isotropic) and b=3000 s/mm^2^. <u>Dataset 4:</u> The dataset from the MASSIVE (Froeling et al., 2017) acquisition was used. In short, we used a subset with 22 b0 volumes and 250 b=3000 s/mm^2^ volumes and an isotropic resolution of 2.5 mm^3^ of one subject.

#### 2.1.2 Image Processing

All images were preprocessed and the diffusion signal was modeled with ExploreDTI v.4.8.6 (Leemans et al., 2009; Leemans and Jones, 2009). Details for each datasets are mentioned below.

<u>Dataset 1:</u> Subject motion, eddy current and susceptibility correction and alignment to structural T1 image was performed by the HCP team according to the HCP minimal process pipeline (Glasser et al., 2013). The fiber orientation distribution (FOD) in each voxel was estimated using CSD (Tournier et al., 2007, 2004) with the recursive calibration method (peak ratio threshold = 0.01) (Tax et al., 2014) with maximum harmonic degree of L_max_=8. For a subset of 10 subjects, we performed additional analysis to test the effects of diffusion modeling. Therefore, we altered only one of the following settings: a) b=2000 s/mm^2^; b) b=1000 s/mm^2^; c) L_max_ =6; d) calibration of the response function based on FA>0.8; e) DTI estimation using REKINDLE.
<u>Dataset 2:</u> Motion-distortion correction and Gaussian anisotropic smoothing was performed with ExploreDTI. The FOD in each voxel was estimated using CSD with recursive calibration (peak ratio threshold = 0.01) and L_max_=8.
<u>Dataset 3:</u> Motion-distortion correction was performed with ExploreDTI. The dataset was resampled to 1 mm isotropic voxel size to increase the level of detail (Dyrby et al., 2014). FODs were estimated using CSD with recursive calibration (peak ratio threshold = 0.01), L_max_=8.
<u>Dataset 4:</u> Dataset 4 was motion, distortion corrected and resampled to 1 mm isotropic voxel size. Similar to the previous datasets, the FOD was estimated using CSD with recursive calibration (peak ratio threshold = 0.01), L_max_=8.

#### 2.1.3 Fiber tractography

For all four datasets, the deterministic FT framework of Jeurissen et al (Jeurissen et al., 2011) was used with parameter settings: FOD threshold = 0.1, angle deviation = 45 degree, step size = 1 mm and minimal tract length = 20 mm. Whole-brain FT was performed with uniform distribution of seed points defined at a 2 mm resolution.

#### 2.1.4 ROI configuration

Automated, large-scale tract selection on the HCP dataset was obtained by atlas based tractography segmentation (Lebel et al., 2008). On the template subject, we defined 4 Boolean ‘AND’ and ‘NOT’ ROIs: 2 axial AND, 1 sagittal NOT and 1 coronal NOT ROIs (fig 1). The 2 axial AND ROIs were placed as follows: the first one was located at the height of approximately half the genu of the corpus callosum and the second ROI was placed 10 slices (equals to 12.5 mm in the HCP dataset) superior. Both AND ROIs included the medial frontal area and excluded the cingulum. A NOT ROI was placed midsagittal to exclude fibers from the corpus callosum. Additionally, one coronal NOT ROI was placed posterior of the frontal lobe and below the corpus callosum to exclude fibers from the inferior fronto-occipital fasciculus (iFOF).

**Figure 1.**
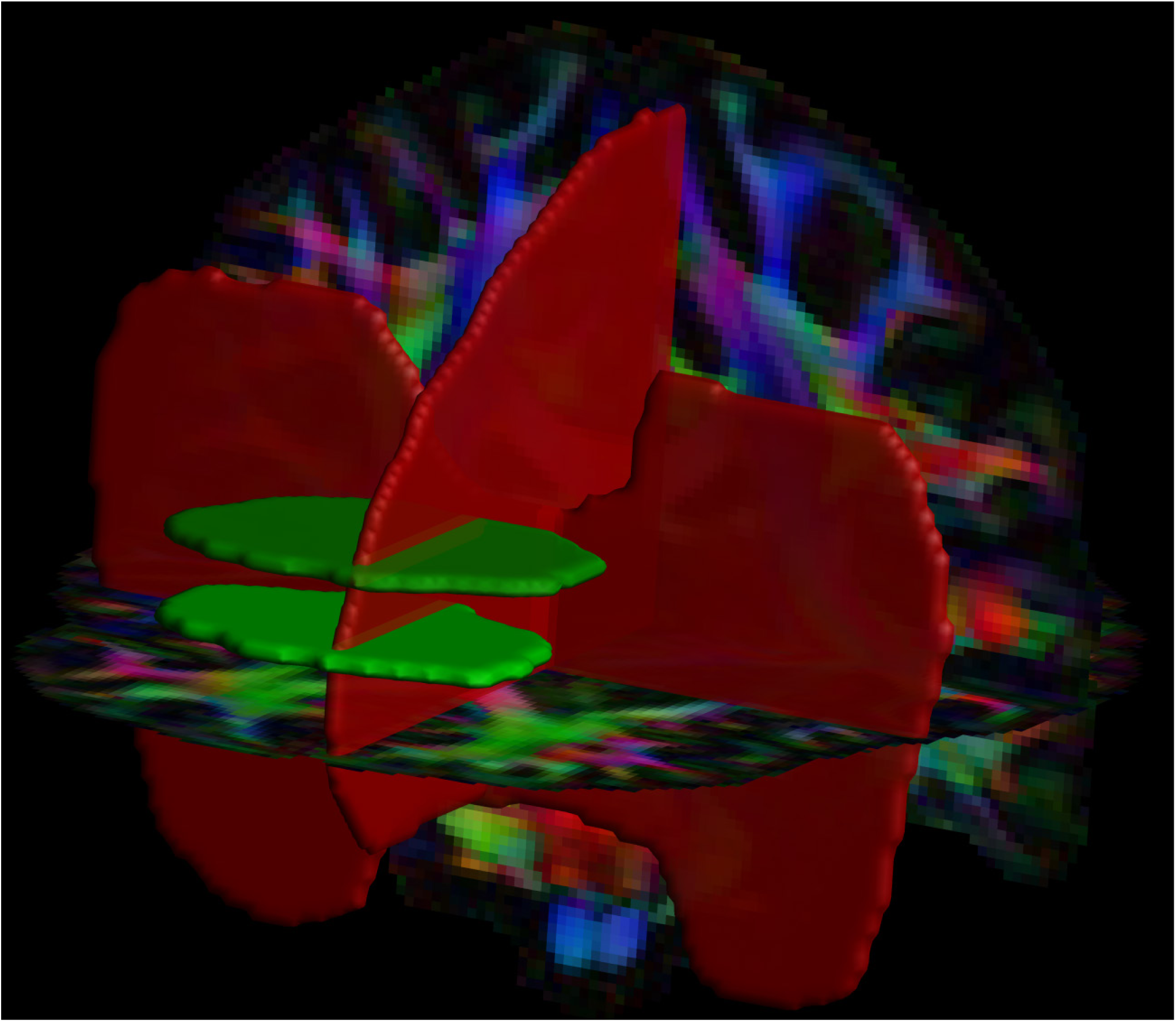
Positions of the two AND (in green) and two NOT (in red) ROIs are shown, which were used for the tract selection in both hemispheres.

#### 2.1.5 Tract consistency

<u>Within cohort:</u> For each of the 409 subjects from the HCP cohort, the visitation mask of the selected tract was normalized to the MNI template (Fonov et al., 2011). The resulting map was thresholded at 1 and binarized. The group composite map displays the fraction of subjects for which the tract mask is presented in each voxel of the MNI space.

<u>Across acquisition protocols:</u> The tractography results of subjects from the four different datasets were visually compared to investigate the consistency of the tract among different acquisition methods.

<u>Across processing settings:</u> We visually checked the effects of different processing settings on the tractography results for 10 subjects of the HCP dataset.

### 2.2 Polarized light microscopy

Polarized light imaging (PLI) is a microscopy method that uses the birefringent properties of myelin sheets to quantify the main fiber orientation in histological sections (Larsen et al., 2007). In current work, we used data previously involved in the investigation of the anterior cingulum (Axer, 2011). One of the studied brains was sliced sufficiently to expose the area of interest. For details on acquisition and processing please see the corresponding study (Axer, 2011). Briefly, a formalin-fixed human brain was macroscopically dissected and a 1.5 cm thick slab of the medio-frontal brain including the anterior cingulum bundle was cut in four blocks. Each of these four blocks were serially sliced and analyzed. The in-plane resolution of the PLI pixels was 64 × 64 μm^2^ with a thickness of 100 μm.

### 2.3 Dissection

Five cerebral hemispheres were anatomically dissected by an experienced neuroanatomist. Dissection was performed after careful examination of the FT reconstructions to guide the separation process in order to limit the loss or damage of the small fascicle of interest. Dissection on the mesial surface started from the cingulate fibers and was continued with the careful investigation of the entire frontal mesial surface.

## 3 Results

### 3.1 Tract description

The complex fiber architecture in the frontal area can be appreciated from the FODs overlaid on the sagittal view in Fig. 2c with the dotted yellow line identifying the interface between regions with locally different dominant fiber populations (“blue” vs. “green” on the principal direction encoded color map in Fig 2 a & b). A medial and frontal view of the bilateral tract configuration is given in Fig. 2 (d) and (e), respectively.

**Figure 2.**
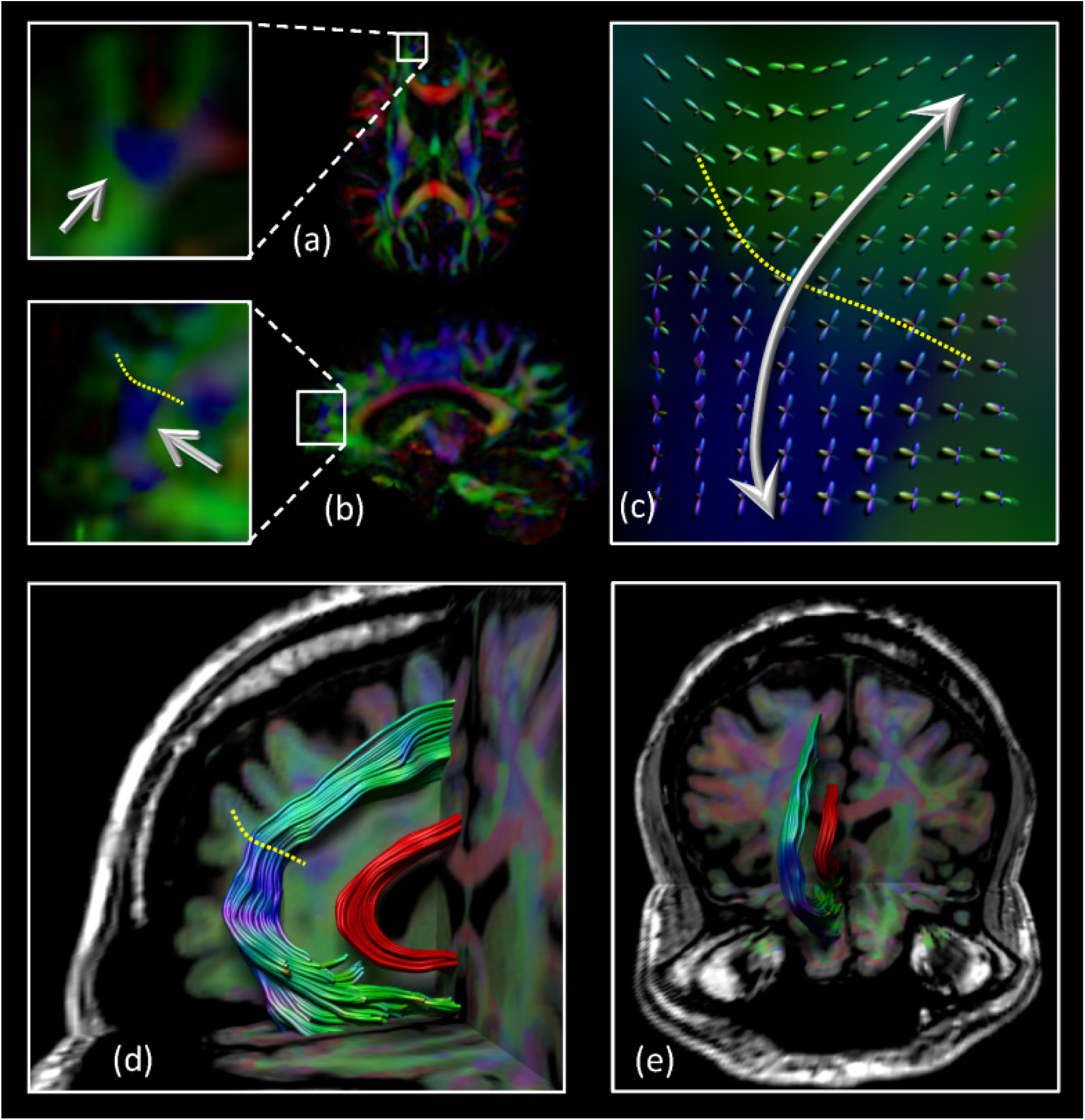
Location of the tract of interest: the axial (a) and sagittal (b) view of part of the tract (blue region indicated by the arrows). The complex fiber architecture in this area can be appreciated from the FODs overlaid on the sagittal view in Fig. 2(c) with the dotted yellow line identifying the interface between regions with different dominant fiber populations (“blue” vs. “green” on the DTI based color map). Medial (d) and coronal (e) views of the fiber bundle configuration. For displaying purposes, only one of the bilateral tracts is shown. The cingulum bundle is shown in red to provide anatomical reference.

The group composite map from the HCP subjects is shown in Fig. 3 where the bilateral tracts follow a similar trajectory as the cingulum, but more frontally (i.e. in front of the cingulate sulcus or within the superior frontal gyrus) and slightly more laterally. The tracts appear to spread from the rostrum of the corpus callosum to the ascending ramus of the cingulate sulcus and are medial to the corona radiata.

**Figure 3.**
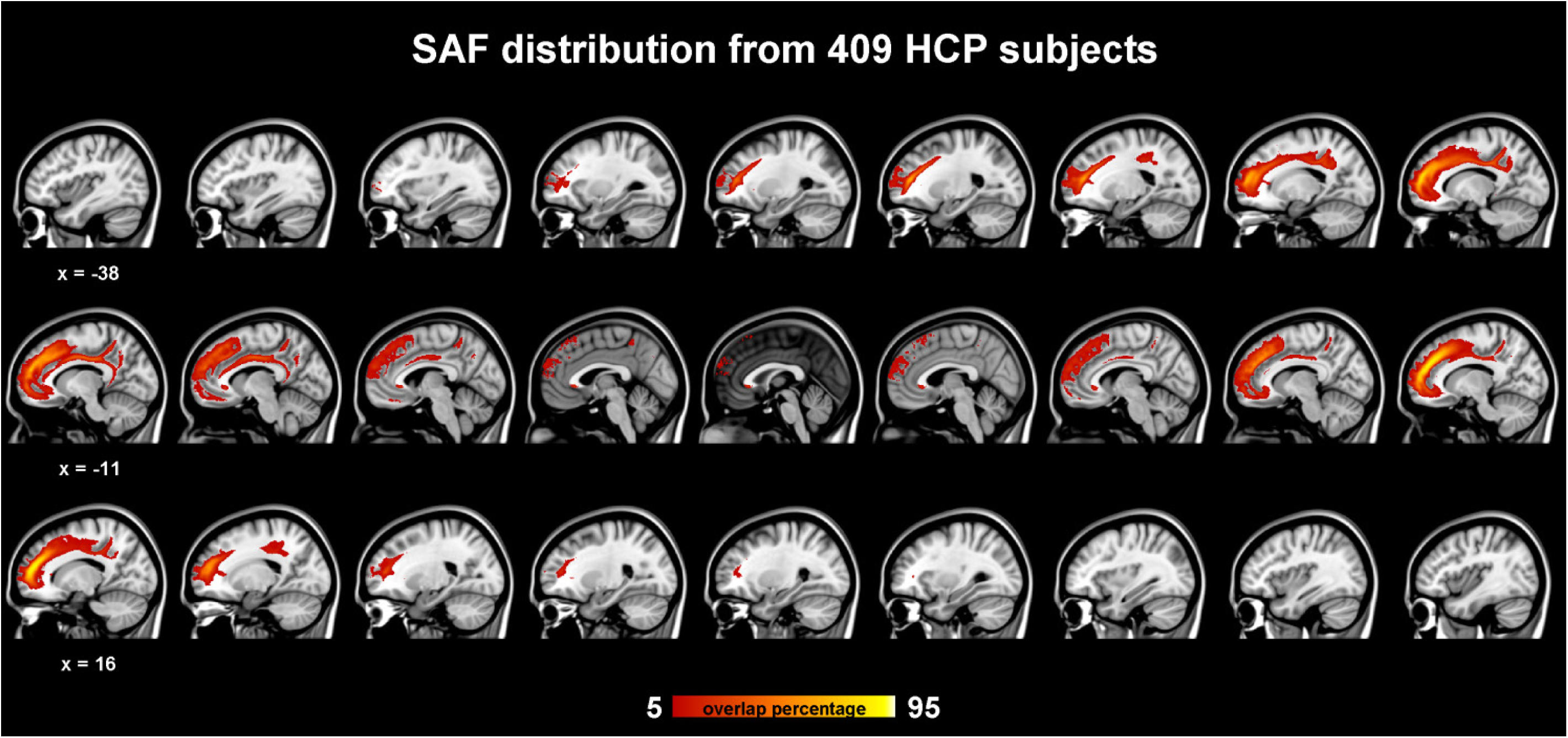
Tract probability map in the 1mm MNI stereotaxic space from 409 subjects of the HCP cohort in sagittal view. The % indicates the fraction of subjects for which the tract is present a given voxel.

### 3.2 Tract consistency

#### 3.2.1 Consistency within cohort/subjects

The percentage overlap map in Fig. 3 shows that the main portion of the tract is present for 90% of the subjects, which is in the same range as other tracts (Thiebaut de Schotten et al., 2011b). The extent of the tracts varied among subjects (see Fig. 4). For some subjects, a very extended and broad structure was found (Fig. 4 bottom row), while there were also subjects in which we found only a few streamlines (Fig. 4 top row). The length of the streamlines varied among subjects, but also within the same subject (e.g. Fig. 4). It is often observed that streamlines stop abruptly due to the absence of a corresponding FOD peak (i.e. in some voxels there appears to be only 1 fiber population presence while the surrounding voxels contain 2 populations).

**Figure 4.**
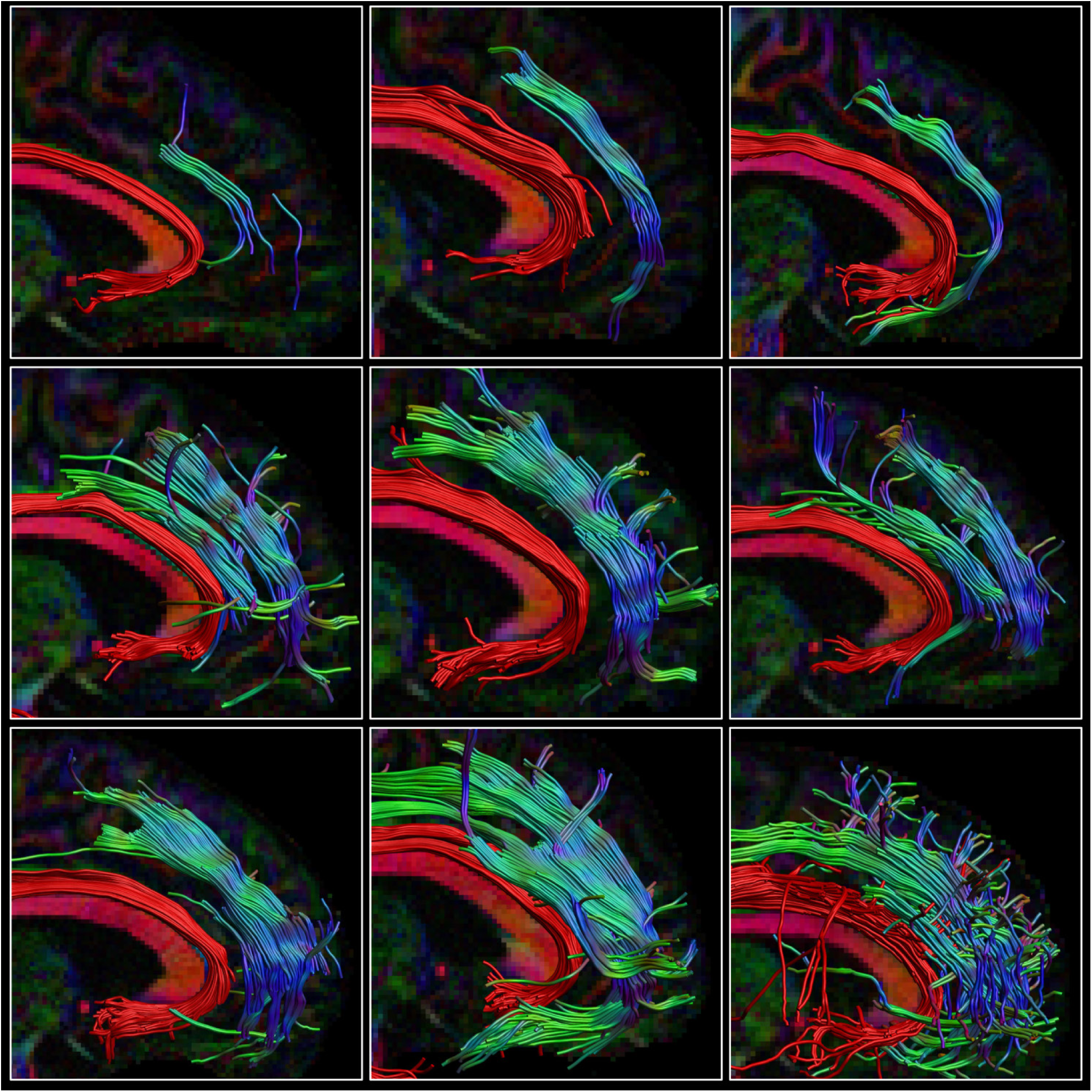
Consistency within the HCP cohort. The right SAF and cingulum is depicted in sagittal view for 9 subjects from the HCP dataset with large variety in extent of the SAF. The cingulum is shown in red to provide anatomical reference. Tracts are plotted on top of the direction encoded color (DEC) FA map.

#### 3.2.2 Consistency across acquisition

Fig. 5 reveals that the tract was present in all cohorts that we analyzed and had a consistent orientation. This emphasizes our observation that the tract is not merely caused by artifacts. While the tract has a consistent orientation, there are differences between acquisition settings, which do have an impact on the extent of the tract. An example is shown in Fig. 6, where the effect of different b-values is depicted. At low b-values shorter and consequently fewer streamlines were found, but CSD based modeling still revealed the tract.

**Figure 5.**
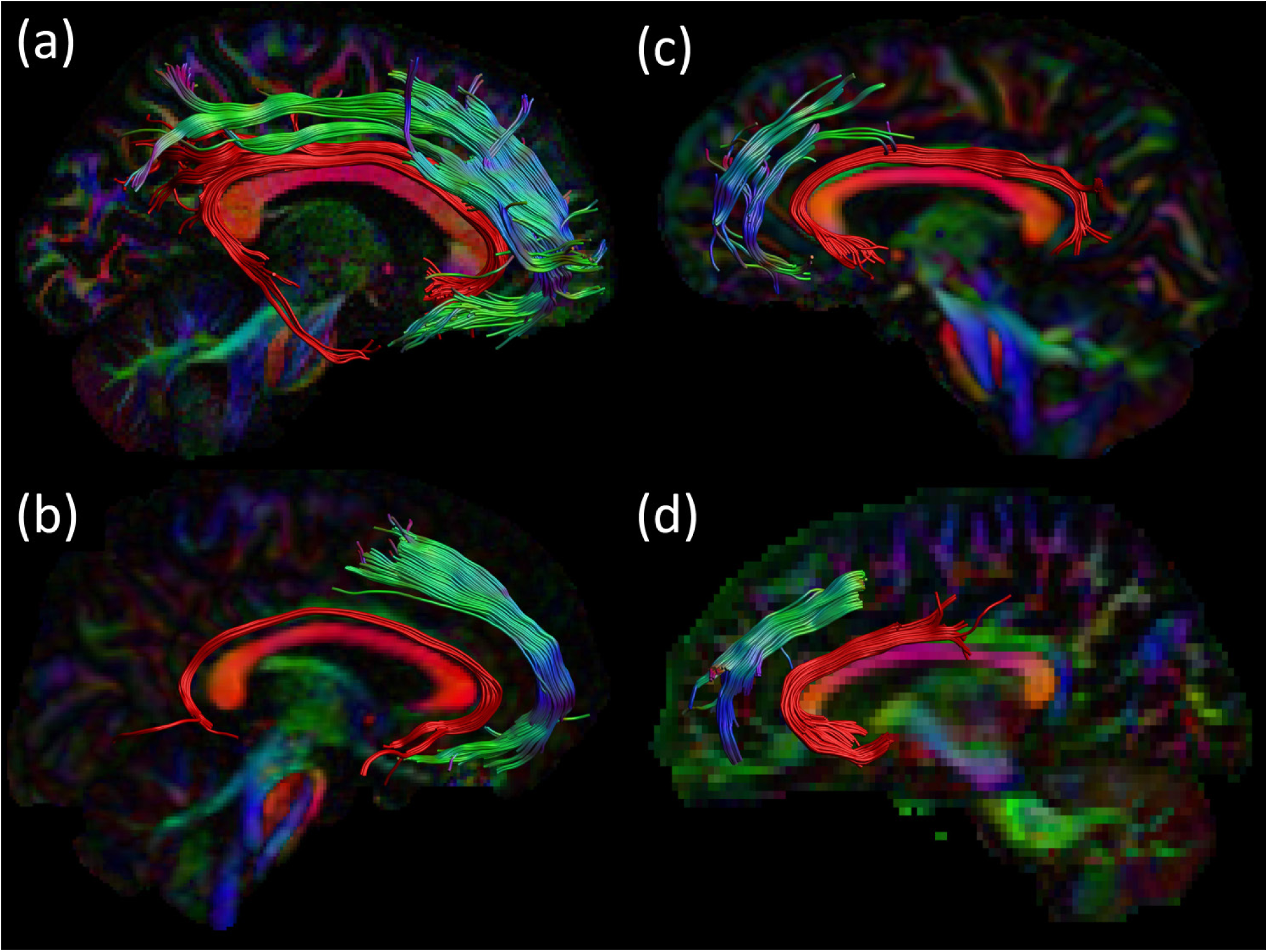
Consistency across acquisition protocols. For all 4 datasets an example of the streamlines is shown in sagittal view: right SAF and right cingulum for (a) HCP and (b) in-house dataset. Left SAF and left cingulum is shown for (c) MASSIVE and (d) ALSPAC datasets. The cingulum is shown in red to provide anatomical reference. Tracts are plotted on top of the DEC-FA map.

**Figure 6.**
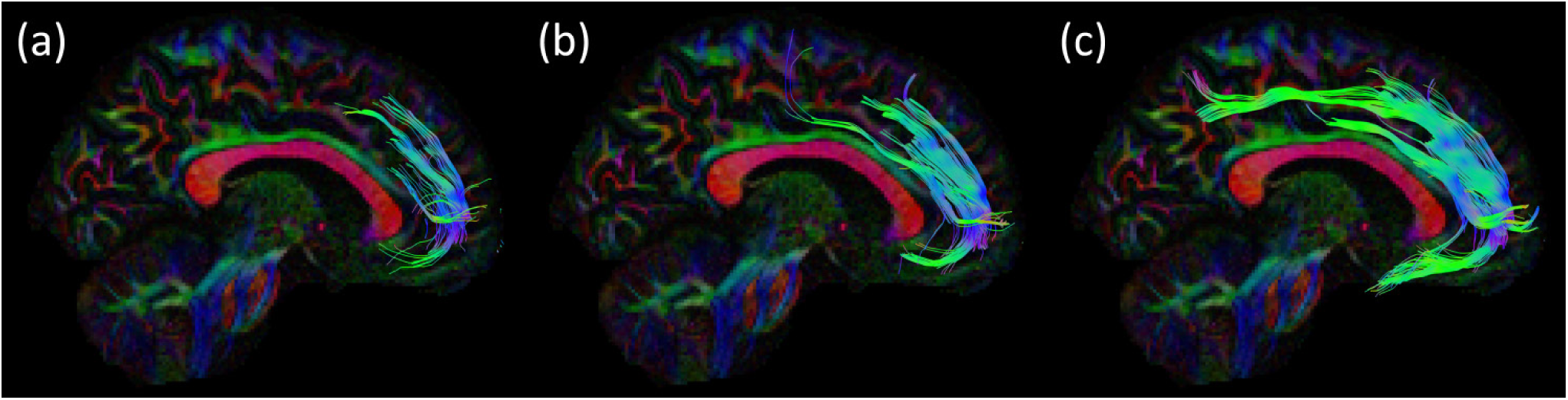
Consistency across acquisition strategies from the same subject of the HCP cohort. The effects of different b-values are shown in sagittal view, other settings are identical: (a) b=1000 s/mm^2^ ; (b) b=2000 s/mm^2^ and (c) b=3000 s/mm^2^. Tracts are plotted on top of the DEC-FA map.

#### 3.2.3 Consistency among processings

In addition, the modeling and processing steps also influenced the extent of the tract (Fig. 7). DTI analysis (Fig. 7a) resulted in either no streamlines at all or in some spurious ones. Alterations of CSD based modeling showed either shorter tracts (L_max_=6, Fig. 7/c) or minor variations (FA calibration, Fig. 7d). Clearly the biggest impact on showing the tract was the modeling of crossing fibers.

**Figure 7.**
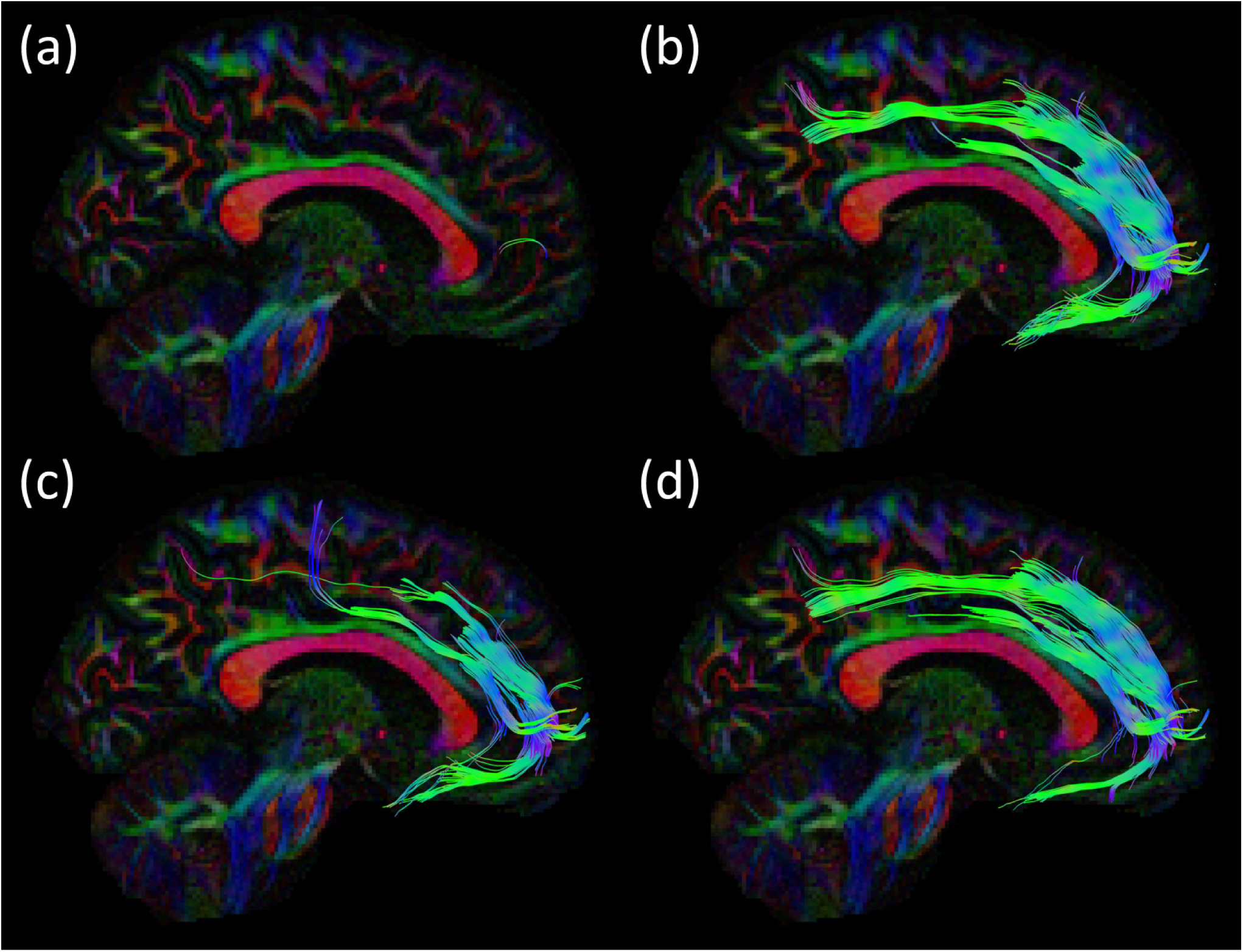
Consistency among modeling and processing solutions from the same subject of the HCP cohort. The effect of modeling or processing on the resulting streamlines are shown in sagittal view: It is clear that the (a) DTI model results in few and spurious tracts, while the CSD models in (b), (c) and (d) produce streamlines, which are plausible with the presented hypothesis. Slight differences in processing can also be observed: (b) recursive calibration and L_max_ =8; (c) recursive calibration and L_max_ =6; and (d) FA calibration and L_max_ =8. Tracts are plotted on top of the DEC-FA map.

### 3.3 PLI

Figure 8 shows the ensemble of the 4 blocks color coded by the main in-plane orientation per voxel. In the region of interest, a variety of colors and therefore locally dominant orientations were present. However, a similar shape (indicated by the white arrows) as with tractography can be appreciated.

**Figure 8.**
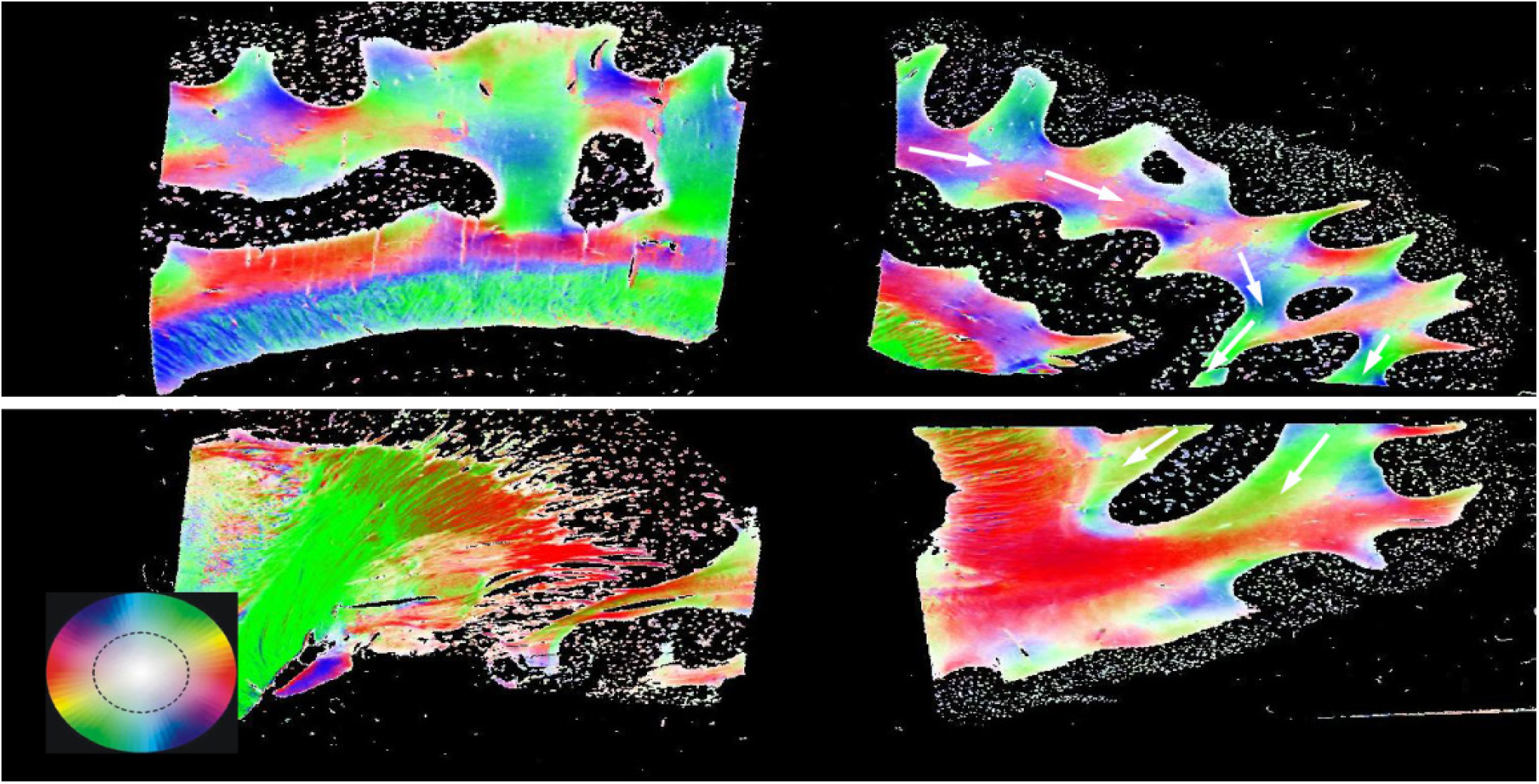
Polarized light imaging (PLI) results. Four separate blocks of the same brain sliced in sagittal plane, where the fiber orientation maps are depicted correspond to the color scheme circle. White arrows are indicating the proposed pathway. The initial motivation of the presented brain was the parcellation of the anterior cingulum bundle according to fiber orientations.

### 3.4 Dissection

With dissection we were able to isolate, in 4 of the 5 specimens, a series of fibers spreading over from the cingulate fibers that seemed to have the same anteroposterior orientation as the tractography tracts. These fibers were located above the cingulum in correspondence with the frontal gyrus (Fig 9). The careful dissection of the fiber complex revealed several different fibers: U-shaped fibers from cingulate, U-shaped fibers belonging to superior frontal gyrus and more lateral fibers from corona radiata (Fig 9). All of them had vertical orientation; therefore no correspondence to the anteroposterior orientation. In one case, the pattern of fibers was not present and interestingly the superior frontal gyrus of that specimen was slim and did not have the usual intragyral horizontal division.

**Figure 9.**
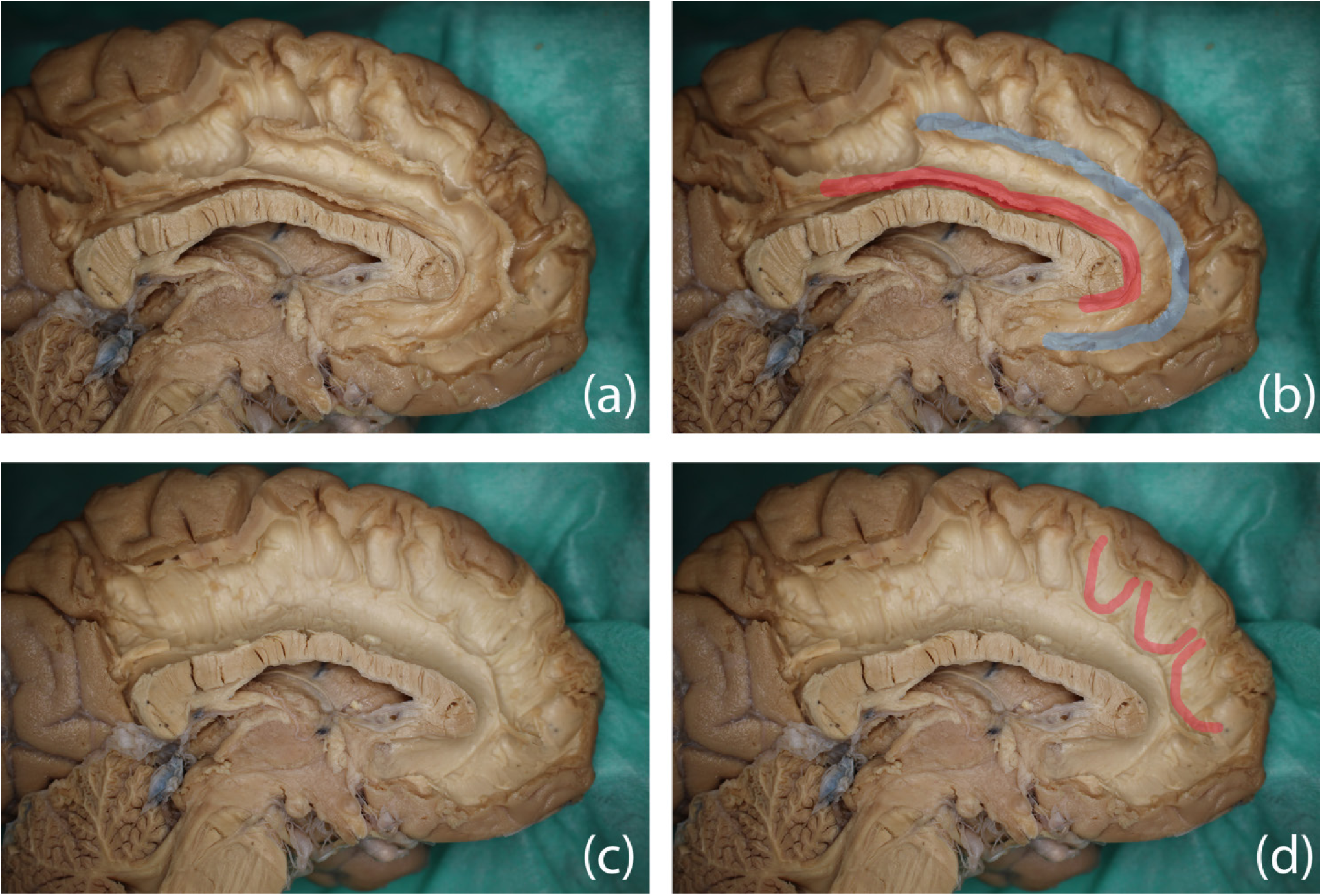
Dissection results in two different hemispheres and color overlays representing the proposed structure. Top row: (a) the original image and (b) possible trajectory of the SAF in blue, the cingulum is in red as the anatomical reference. Bottom row: (c) original image and (d) several U-shaped fibers, which can form the SAF.

**Figure 10.**
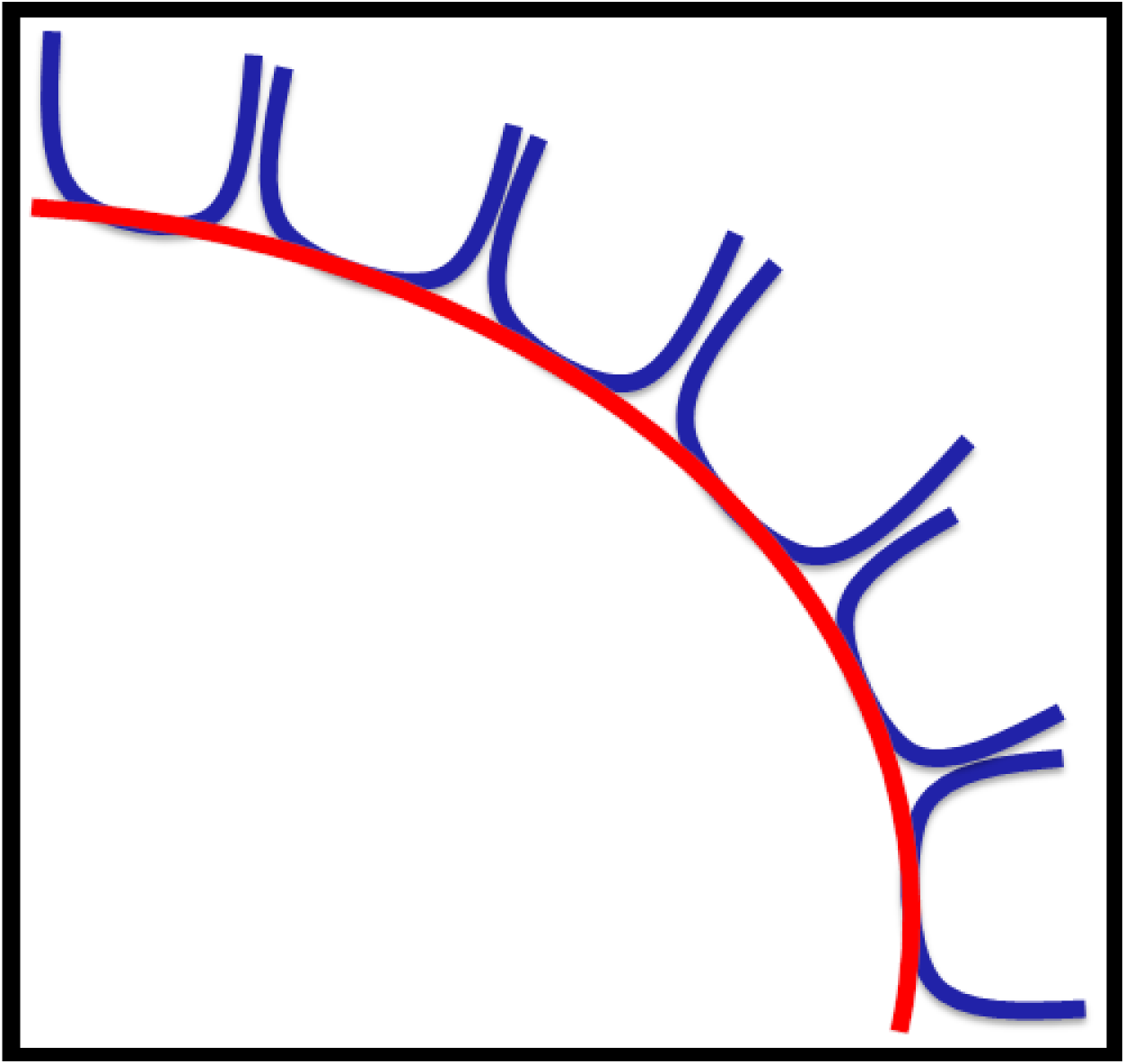
Hypothesis on how consecutive U-shaped fibers (in blue) can form a long pathway (in red).

## 4 Discussion

By taking advantage of state-of-the-art diffusion MRI methodology, we have identified a consistent bilateral tract via fiber tractography in the prefrontal lobe, which has not been described before (e.g., see previous work on FT (Catani et al., 2012; Makris et al., 2005; Thiebaut de Schotten et al., 2011b). Reproducibility across multiple subjects and different data cohorts and acquisition settings boosted our confidence that this finding is not based on imaging artifacts.

FT shows a consistent tract, however, the presence of a long association fiber is not fully supported by dissection which necessitates hypothesizing about why a pathway is found. A plausible explanation is that the series of consecutive U-shaped fibers that connect adjacent gyri emerge to form a long pathway (see Fig 8). This hypothesis is supported by work of Maldonado (Maldonado et al., 2012) where they propose that the dorsal component of SLF is primarily consisted of U-shaped fibers.

While the agreement between dissection and FT is partial, it is important to describe the structure and recognize its occurrence. First, as this is a consistent finding, other researchers will likely discover and investigate the presence of this FT structure. Second, investigation of the structural properties of the sub-compartments can improve the understanding of the U-fibers. Third, this adds to the discussion of the value of FT, thereby raising additional awareness of the pitfalls of FT (Jones and Cercignani, 2010; O’Donnell and Pasternak, 2015; Parker et al., 2013) and the concerns in the field of connectomics (Fornito et al., 2013; Hagmann et al., 2010; Zalesky et al., 2016).

In the following sections, we will discuss that our FT results are plausible from the point of dMRI-based tractography, place our results in the context of other FT/dissection studies, provide suggestions for future investigations, and stimulate the debate about the validity of dMRI-based FT.

### 4.1 Diffusion MRI

Recent developments in acquisition and data processing boosted the reliability of dMRI and increased the inherently low accuracy of mapping WM pathways with FT. The investigations in solving crossing fibers were imperative, as the reported structure would not or hardly be detected since the main diffusion direction here is along the forceps (see Fig 2).

However, one of the pitfalls of dMRI is that it is an indirect measure of the underlying WM pathways and reflects the net displacement of water along all structures within a large voxel. Different structural architectures can lead to the same diffusion profile (Jbabdi and Johansen-Berg, 2011), making it hard to unambiguously reconstruct the intrinsic fiber arrangement.

#### 4.1.1 Tract consistency

The tractography result shows a high similarity of 90% for the core of the tract, which is outstanding given the fact that WM fiber bundles vary in their size and position, and this is a small and inferior bundle. Compared to other larger bundles, the similarity is substantial related to previously reported overlap of >90% (cingulum), >90% (core CC), >75% (CC), and >75% (inferior fronto-occipital fasciculus) (Thiebaut de Schotten et al., 2011b).

The presence of the tract among different data cohorts and acquisition settings is indicative that the results are not merely based on acquisition artifacts and that the structure can be found in all datasets using current high angular resolution acquisition. As FODs are sharper, more and longer streamlines are found at higher b-values as this aid the resolving power of the different fiber populations within a voxel.

Although we can show the tract in case of all the subjects, finding the complete tract is not trivial. The tract can be interrupted, for example when the FOD only contains the main peak of the crossing fibers and does not contain the minor second peak. While we were able to determine the main part of the tract, the terminations are hard to obtain due to error propagation along the tracts. Due to the underlying complexities in finding the complete tract, one should currently refrain from relating tract properties (e.g. length and size) to anatomical features in group studies.

U-shaped fibers connect neighboring gyri throughout the brain and form an area of investigation (Rojkova et al., 2016; Zhang et al., 2014). However, performing fiber tractography on U-fibers is difficult due to the high curvature within these fibers and moreover different FT settings are needed, e.g. larger curvature setting. Even after adjustments of these settings for a few datasets, we were not able to clearly display the U-fibers in the frontal region to determine if the pathway of the U-fibers and the described pathway do overlap. Future studies, especially at high resolution, might alleviate this question.

### 4.2 Validation

Validation with PLI and dissection is showing dissonant results. PLI shows an overall pathway that can resemble the presented structure; however, it is not clearly one consistent pathway. A drawback of PLI is that it cannot distinguish crossing fibers, making it hard to be unambiguous about the exact pathway. Fiber dissection did not confirm association fibers, but points toward a series of fibers. A critical note on dissection is the limitation that small fibers are hard to dissect, especially when the majority of the fibers are crossing. Small fibers might survive better using a different dissection technique, i.e. Klingler’s dissection (Agrawal et al., 2011; Klingler, 1935). As stated before, the hypothesis that the FT-based tract is formed by a succession of short U-shaped fibers is also posted by Maldonado, who dissected short U-fibers at the location of the long association fibers of the SLF. In summary, the invasive methods did not verify the SAF clearly, possibly due to the limitations of these methods or to the presence of the bundle, however it could not provide exclusionary reasons either.

### 4.3 Complexity of fiber bundles

Interaction between tractography and dissection methods drives both the validation of tracts, but also the discovery of true fiber bundles, thereby increasing our understanding of the design of the brain’s architecture (De Benedictis et al., 2016; Hau et al., 2017; Meola et al., 2015; Wu et al., 2016; Yeatman et al., 2014). Over the years more and more (parts of) WM bundles were revealed using FT that previously were undetectable with the formerly existing techniques (Jbabdi and Johansen-Berg, 2011), such as the lateral projections of the corpus callosum (CC), IFOF or the Aslant fiber bundle (Thiebaut de Schotten et al., 2012). However, the validity of some of these tracts is still debated. Yeatman et al. (Yeatman et al., 2014) used tractography to rediscover the vertical occipital fasciculus (VOF); a bundle that caused controversies among neuroanatomists in the 19^th^ century. These tractography findings were further confirmed via dissection by Wu et al (Wang et al., 2016).

Other researchers have also proposed new or redefined WM structures. Track density imaging (TDI) on 7T dMRI (Calamante et al., 2012, 2011, 2010) recently demonstrated finer details of thalamocortical connections (Choi et al., 2018) and revealed the fiber of the septum pellucidum area (Cho et al., 2015). In both studies the super-resolution ability of the voxelwise fiber count was used to generate images with high anatomical contrast and therefore exposed the aforementioned bundles. However, due to the noise sensitivity of the technique, the validity of structures revealed by TDI remains an open question (Dhollander et al., 2014, 2012).

In addition to new bundles, several studies have shown subcompartments of known WM pathways, which are validated by histology. These multi-component bundles such as uncinate fasciculus (Hau et al., 2017), inferior fronto-occipital fascicle (Sarubbo et al., 2013) and SLF (Kamali et al., 2014; Makris et al., 2005) consist of a complexity of pathways that together form a united bundle. De Benedictis et al. (De Benedictis et al., 2016) used a microdissection approach to reveal and validate the presence of both homotopic as well as heterotopic fibers in the anterior half of the CC.

Interestingly, the cingulum, which has a similar shape and is in close approximation of the presented tract, is also a multi-component bundle. The cingulum bundle consists of many short fibers as well as longer fibers that together have many different connections (Golding and ALSPAC Study Team, 2004). The cingulum is usually depicted as a continuous structure using fiber tractography although thorough research is showing a division in at least three subparts, each with their own distinct diffusion metrics (D. K. Jones et al., 2013; Wu et al., 2016). Given the similarity with the cingulum and the knowledge that many bundles have complex and multicomponent fiber organization, it seems plausible that the newly described frontal tract also has a multicomponent organization.

### 4.4 Future directions

The study describes a provoking fiber tractography finding that potentially has implications for the FT-community and for neuroscience. However, further research is needed for better understanding of the origin of this structure. FT related topics can lead to the investigation of this area at very high spatial resolution for a finer understanding of the complex structures (Jeurissen et al., 2013) as the prevalence of fiber crossings is increasing with the spatial resolution (Schilling et al., 2017). This would ideally be combined with careful dissection. Another route is inspecting this tract with even more advanced FT methods that for example incorporate anatomical or contextual information (Daducci et al., 2016; Smith et al., 2012).

We observed a large variability in the cross-sectional area of the tract and future analysis of the structure should entail examining common measures such as microstructure, shape, demographics, and pathological changes. It is important to point out that the frontal lobe is generally challenging to investigate by most MRI methods, because of the susceptibility induced distortions, which are further emphasized by EPI sequences caused by the air-tissue interfaces of the sinuses. The dedicated effort to correct for such acquisition artifacts by the HCP team (Andersson and Sotiropoulos, 2016, 2015; Sotiropoulos et al., 2013) made it possible to investigate the tract in a large cohort.

### 4.5 Conclusions

To summarize, we presented comprehensive evidence of a white matter bundle with dMRI tractography, while it is also probable that this does not exclusively resemble a long association fiber. We showed a consistent FT finding that is present within several cohorts, among different acquisition settings and processing strategies. However, we are uncertain about the true underlying anatomical structure. The current, most reasonable explanation of the presence of the FT tract is that it follows a route laid out by shorter (U-shaped) fibers, similar to a highway, where traffic enters and leaves the highway and travels only on a section of the total highway. So far, DTI gave a description of highway-like structures, while advanced modeling subsequently allows us to see less dominant pathways as well.

